# PCA-based source-space contrast maps reveal psychologically meaningful individual differences in continuous MEG activity

**DOI:** 10.1101/835884

**Authors:** Erkka Heinilä, Aapo Hyvärinen, Tapani Ristaniemi, Lauri Parkkonen, Tiina Parviainen

**Author notes:** Corresponding author. Faculty of Information Technology, University of Jyväskylä, Finland, PO Box 35 FI-40014 University of Jyväskylä.

## Abstract

Within the field of neuroimaging, there has been an increasing trend towards studying brain activity in naturalistic conditions, and it is possible to robustly estimate networks of on-going oscillatory activity in the brain. However, not many studies have focused on differences between individuals in on-going brain activity that would be associable to psychological or behavioral characteristics. Existing standard methods can perform well at single-participant level, but generalizing the methodology across many participants is challenging due to individual differences of brains. As an example of a clinically relevant, naturalistic condition we consider here mindfulness. Trait mindfulness, as well as a mindfulness-based intervention cultivating focused attention, is often associated with benefits for psychological health. Therefore, the manner in which the brain engages in focused attention vs. mind wandering is likely to associate with individual differences in psycho–behavioral tendencies.

We recorded MEG from 29 participants both in a state of focused attention and in a state of simulated mind wandering. We used Principal Component Analysis to decompose spatial average activation maps of focused attention contrasted with two different mind wandering states. The first principal component, which reflected differential engagement of bilateral parietal areas during focused attention vs. mind wandering, was associated with behavioral characteristics of inhibition, anxiousness and depression, as measured by standard questionnaires. We demonstrated that such decomposition of time-averaged contrast maps can overcome some of the challenges in methods based on concatenated data, especially from the perspective of behaviorally and clinically relevant characteristics in the ongoing brain oscillatory activity.

**Highlights:**

- We present a specific method to analyse/establish associations between brain oscillations and behavioral characteristics.
- We found that activity levels in parietal areas during mind wandering compared to focused attention were associated with the behavioral trait of inhibition and anxiety.

## 1. Introduction

Studies using functional magnetic resonance imaging (fMRI) and recently also magnetoencephalography (MEG) have shown that even with no specific task, brain exhibits robust patterns of activation that are consistent over individuals (Nugent et al., 2015; Ramkumar et al., 2012). However, the lack of a specific task makes it difficult to use standard methods that are designed for correlating activation patterns with known task or stimulation patterns. Thus, unsupervised methods such as Independent Component Analysis (ICA) have become popular in this context. After a breakthrough fMRI study (Beckmann et al., 2005) where many such resting-state networks were identified in a data-driven manner using ICA, similar networks that are consistent over subjects have also been identified in brain electrophysiology using MEG (Brookes et al., 2011).

While it is important to study the commonalities over individuals, for better understanding the relevance of resting brain dynamics in individual experience and behavioral tendencies, we also need to explore how individual brains differ in these task-free conditions. Indeed, with an increase in sample size and use of more sophisticated data analysis methods, it is becoming feasible to extract features in brain activation patterns that associate with individual trait differences in cognition or behavior (Mason et al., 2007). For example, the frequency distribution of spectral content of resting state electrophysiological activity in the brain shows stability over individuals (Näpflin et al., 2007) and reflects the level of cognitive load (Haegens et al., 2014).

However, there are very few investigations that have focused on the link between resting-state oscillatory dynamics and psychological traits within a typical healthy sample of participants. Interestingly, MEG was recently used to show that increased beta-band modulation to anxiety-evoking images differs between participants who profile at the opposite ends in the dimension of tendency to be exploratory vs. tendency to be cautious (Yamano et al., 2016). In that study, the differences were shown at the group level by classical hypothesis-driven t-statistics on activation in response to stimulation. Yet, the results indicate that there are indeed features in the MEG oscillations that may reflect, in a fundamental way, how we approach the environment, especially in emotionally loaded contexts. Building on this assumption, data-driven characterization of brain oscillations would enable comparisons of general resting-state dynamics that underlie psychological differences in human behavior.

However, the multi-participant context requires additional care regarding the decomposition of data into components. In order to achieve a meaningful interpretation, the resulting brain oscillation components for each participant should be comparable to each other. Running ICA separately for each participant in a group results in a set of components for each participant with no simple way of matching them (but see Esposito et al., 2005). Organizing original data of each participant in some common structure, by, for example, spatial or temporal concatenation, allows to run ICA only once yielding components in common space in one go, and then the participant-specific components can be extracted using back-projection. This has become a popular choice to do ICA on fMRI at group level (Calhoun et al., 2009). While it has been used successfully for example to find connectivity differences between healthy participants and participants with major depressive disorder (Nugent et al., 2015), it also has drawbacks. As the datasets are concatenated before ICA is run, the resulting components are all constrained with the same ICA mixing matrix. This means that the independent processes that ICA models, are assumed to be (either spatially or temporally) consistent over participants. It is not completely clear what the implications of this assumption are, for example, in the case of inconsistency of measurement errors or other differences between participants, even though the matter has been investigated to some extent (Allen et al., 2012). In addition, some components may represent processes that are based only on a subset of participants. For a study on group-level inferences, aiming to examine the link between brain activation and individual differences, this poses a problem: how to know if a back-projected component of a participant is simply noise (if the component was in reality reflecting other subjects’ activities) or represents a participant-specific modulation of the common independent process. Finally, even if the estimation went fine, ICA typically results in a lot of components. It is a common challenge to determine which components are selected for further analysis. Without valid a priori information in the component selection, this can lead to selective reporting or loss of power due to multiple comparisons.

Mindfulness meditation provides an interesting testbed to develop and test methods for continuous data analysis, as it represents a distinct, yet ‘trigger-free’ condition. In the context of meditation, practicing focused attention is often associated with many health benefits, such as optimal emotion regulation, cognitive control, stress reduction and feeling of fuller life. It can thus be hypothesized that individual differences in the neural signatures of focused attention would correlate with behavioral tendencies and personality characteristics that link with psychological wellbeing.

We recorded MEG in a paradigm where a sample of 29 participants were asked to perform focused-attention meditation task (FA) and two different control conditions: reflection on anxious thoughts (AT) and self-centered future planning (FP). These latter conditions served as a simulation for mind wandering, which is generally considered as an opposite state of mind to mindful focused attention. Thus, importantly, these two control conditions could be used as a meaningful baseline to extract the essential features reflecting sustained attention during mindfulness meditation.

For analysis of this data, to alleviate the problems discussed earlier, we devised an alternative approach based on temporal averaging and contrasting instead of temporal concatenation. We first projected sensor-level time courses to source space, and then computed amplitude envelopes of band-pass filtered data from these with Hilbert’s transform. We focused on alpha oscillations as they have good signal-to-noise ratio and they have been shown to be modulated by selective attention (Foxe & Snyder, 2011). Instead of using ICA on concatenated time courses, we computed average amplitudes for each brain voxel and for each task, and then did pairwise subtractions to get spatial contrast maps, that is, the difference of amplitudes in each of the voxels. This was done separately for each subject and for all three pairs of tasks (FA-AT, FA-FP and FP-AT). Feeding these spatial contrast maps into Principal Component Analysis (PCA) resulted in one robust component, which explained most of the variance across subjects, for all three task pairs. We then used regression to explore the significance of the state-related cortical differences, represented by the principal components, for psychological traits, measured by temperament and personality questionnaires.

The proposed pipeline mitigated many of the problems discussed above for analysing the connection between resting-state dynamics in MEG data and specific individual (trait or state) characteristics. Because the within-participant variation is averaged out, the decomposition is not based on processes on individual participant level, but instead on the differences between participants, making interpretation straightforward. Even though intricate information on the within-participant fluctuations is lost, for the task of exploring individual differences it is meaningful to focus on subspaces that reflect the variation between participants. The use of spatial contrast maps further increased the specificity of the method, as the subtracted contrast tasks essentially acted as a baseline. This removed both within- and between-participant variability not related to tasks. These steps together made the final collection of components relatively small, and thus less statistical power is lost in the component selection. This is especially important with a relatively low sample of participants, which is typical in neuroimaging studies.

## 2. Materials and methods

### 2.1. Participants and data acquisition

The project was run as a collaboration between Aalto University, University of Jyväskylä and University of Helsinki. 29 people (aged 21–48 years) took part in the initial data gathering phase. Participants had no history of neurological disorders, head trauma or substance abuse, and had normal or corrected to normal vision. Twelve participants had no previous meditation experience, while other participants had meditation experience ranging from 0.5 to 10 years. For each of the participants, we collected two 30-minute recordings of magnetoencephalography (MEG) with Elekta Neuromag TRIUX system (MEGIN Oy, Helsinki, Finland). Each recording session included two 2-minute resting-state blocks at the beginning and at the end of the sessions. In between there were three different 8-minute tasks organized in 2-minute blocks in counterbalanced order (see Fig 1.)

**Fig 1.**
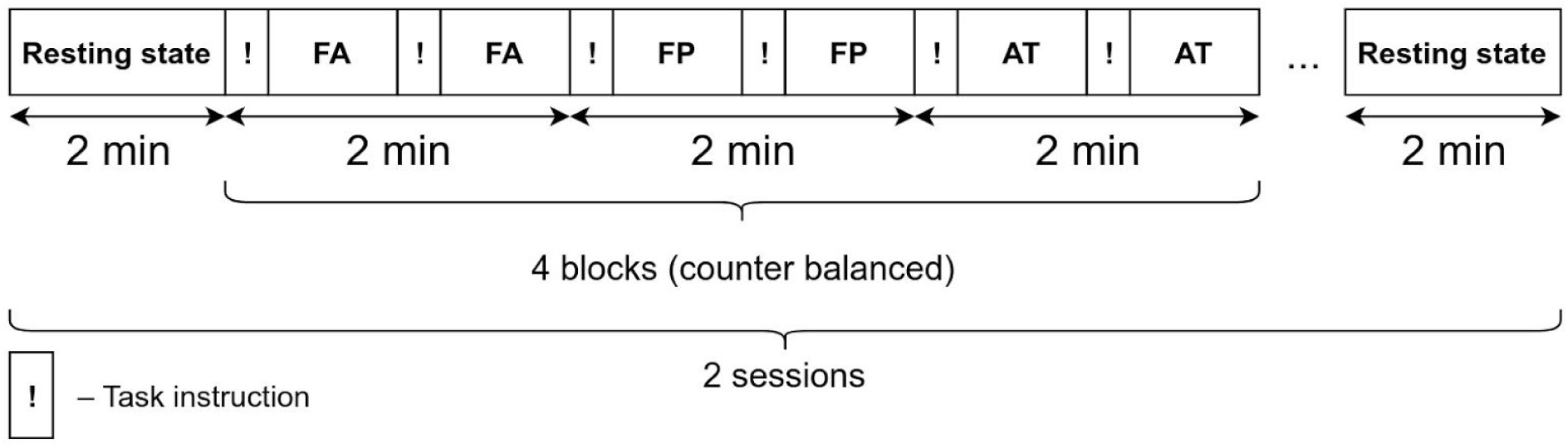
Experiment design. Tasks of focused attention (FA), future planning (FP) and anxious thoughts (AT) are carried out in counterbalanced order. Each 1-minute block contains a brief instruction in the beginning (!).

The three tasks included were focused attention on breathing (FA), future planning (FP) and anxious thoughts induced by reflection on emotional pictures (AT). In all tasks, participants were instructed to sit still, fix gaze on a crosshair in front of them, and perform the task. Before each 1-minute miniblock, an instruction picture was shown with a text: “Focus on your breathing” for the FA task, “make plans related to the picture” for the FP task, and “place yourself or someone close to you in this situation” for the AT task. Pictures were selected beforehand by the participant from International Anxious Picture System (IAPS) -database to maximize the affective experience for each participant individually.

### 2.2. Trait questionnaires

Psychological and behavioral traits were characterized by using standardized questionnaires. Based on earlier studies (Lyyra and Parviainen, 2018; Schneider et al., 2018; Yamano et al., 2016) we hypothesized that traits along the dimension of behavioral inhibition and behavioral approach as well as anxiety would most likely be captured in the brain dynamics, and therefore focused on questionnaires in this domain. These included questionnaires related to anxiety, depression and behavioral inhibition or activation. BIS/BAS is a self-report questionnaire designed to measure two motivational systems: behavioral inhibition system and the behavioral activation system (Carver and L. White, 1994). BDI measures characteristic attitudes and symptoms of depression (Beck et al., 1961). BAI is used for measuring the severity of anxiety in children and adults (Beck et al., 1988).

### 2.3. Preprocessing of trait data

To lower the dimension of possibly collinear questionnaire results, we ran PCA on the data originally containing variables for BDI, BAI, BIS, BAS Reward Responsiveness, BAS Fun Seeking, and BAS Drive questionnaires from all participants.

### 2.4. Preprocessing of MEG data

MEG was used to record magnetic fields from 306 channels arranged in a helmet around the head of a participant. Recording was done with 1000 Hz sampling frequency, yielding 306 time series. Before the recording, the shape of participant’s scalp was digitized and the five coils attached to the head were localized with respect to anatomical landmarks (nasion, preauricular points). During the MEG measurement, weak current with specified frequency was applied to the coils so that the location of sensors with respect to the helmet could be determined.

The recordings were preprocessed with the signal-space separation method (MaxFilter 3.0; MEGIN Oy, Helsinki, Finland) to suppress magnetic interference originating from outside the helmet. As the head location in the helmet can vary from participant to participant, and can change during the recording, MaxFilter’s utilities for movement compensation and translation to the same virtual head location were used. Common artifacts such as eye blinks and heartbeats were extracted and removed semi-automatically using ICA implemented in MNE-Python (Gramfort et al., 2014). One participant had to be completely dropped from further analysis because of low signal-to-noise ratio.

To enhance interpretability, data were further transformed from sensor space to source space, that is, a model of neural currents within the brain, using the following procedure. For each participant, the digitized points from the measurement were fitted to average head model from the freesurfer software package. As head size can vary between participants, we first uniformly scaled (scaling factors varied between 0.85 to 1.0) the average head for each participant separately so that the fit would be as good as possible. Both the scaling and fitting were done using MNE-Python’s coregistration utility.

Within the head model, we used a volumetric source dipole arrangement. Instead of using the same spacing for each participant, we first constructed source space for the original freesurfer model with 8mm spacing, and then scaled the source space to match individual participants. This yielded one-to-one correspondence between voxels of the head models, making comparisons between participants simple later on.

The actual inverse problem was then solved using Minimum Norm Estimate (MNE) -method in combination with noise normalization by dynamic Statistical Parametric Mapping (dSPM).

### 2.5. Data Analysis

The overall dataset consisted of two independent 8-minute recordings for each of the continuous tasks (FA, FP, AT) from 28 participants. Solving the inverse problem yielded time series for each voxel in the head model. We band-pass filtered the signals to a range of frequencies known as the alpha band (7-14Hz), and then, using Hilbert transform, computed amplitude envelopes over time. For each task, participant and voxel separately, we computed the average over all time points of the 16-minute concatenated envelope, resulting in one spatial map for each task and each participant. While we were primarily interested in the differences between the tasks, association of these task-specific spatial maps to behavioral characteristics was investigated as a preliminary step.

Next, we computed spatial contrast maps. This was done for all three task pairs (FA-FP, FA-AT, FP-AT) as a simple subtraction of the task-specific spatial maps, resulting in n_voxels-dimensional (where n_voxels is the amount of voxels) vector for each task pair and each participant. The following steps were done for each task pair separately, i.e. analysing variability over the 28 participants for each task pair.

To focus on the most relevant interactions between individual trait characteristics and task-related brain activity, we conducted PCA to extract subspaces containing most of the variance. To evaluate the robustness of PCA in our context, we ran a variant of icasso (Himberg et al., 2004) to reveal how sensitive the resulting components were to the presence of individual participants.

The icasso-based evaluation procedure went as follows. For each participant, we picked the spatial contrast maps for all except this particular participant, and stacked them to a (n_participants - 1) * n_voxels matrix. This matrix was then decomposed to three principal components and their corresponding mixing weights (spatial weight maps) that explain most of the variance. We used hierarchical clustering to cluster all the spatial weight maps from all runs pooled together. The compactness of clusters, indicating robustness to individual participants being present or not, was inspected visually from a dendrogram. After evaluation, we ran the PCA once more with all the participants, and used that decomposition in the subsequent steps. We additionally ran ICA in the subspace spanned by the three principal components to see if using higher order information can complement the analysis.

Finally, we used linear regression to model the relationship between the first principal component of the spatial contrast maps and the first principal component of the questionnaire data. Statistical significance of the regression coefficient was evaluated with a standard t-test.

## 3. Results

### 3.1. Behavioral inhibition — approach -component

We used PCA to decompose the data from questionnaires to three components. The mixing weights are shown in Fig 2. The first component explained 50% of the data. As the component seemed to capture positive correlations between the BDI, BAI, and BIS questionnaires, but also negative correlation to BAS Drive, we call this behavioral inhibition – approach -component (BIA). The other two components (see Fig A.1) explained both less than 20% of the variance, and were not as intuitively understandable, so they were left out for the subsequent analysis.

**Fig 2.**
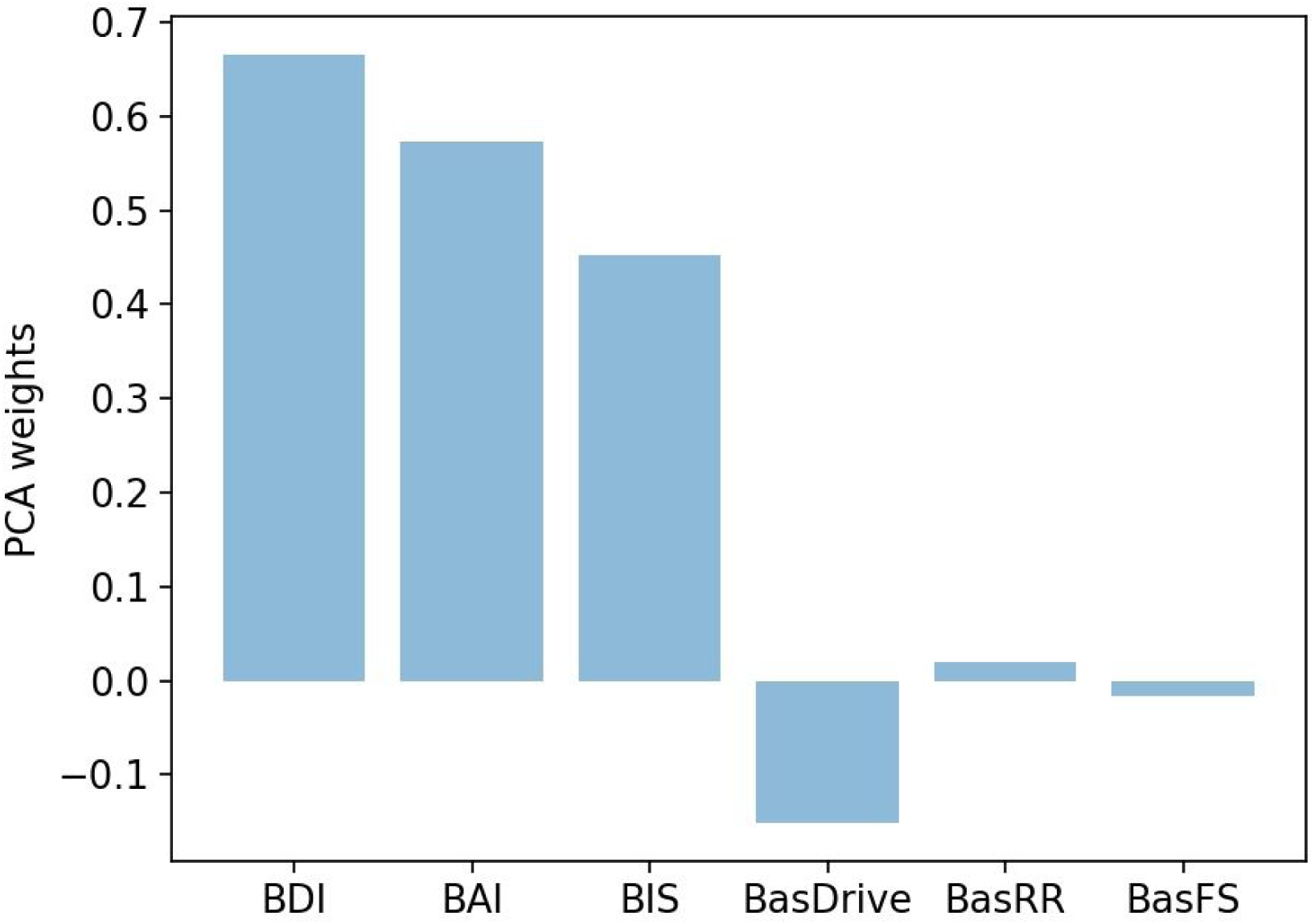
Mixing weights of the first PCA component computed from questionnaire data (behavioral inhibition – approach component.)

### 3.2. Spatial contrast maps

Using the procedure detailed in the methods section we created spatial contrast maps from MEG data for all participants and for all task pairs. Averages computed over participants for each task pair (FA-FP, FA-AT, FP-AT) are shown in Fig 3. In the figure, the red color means that the amplitude of alpha oscillations during the second task (FP in FA-FP pair) is higher than the amplitude of alpha oscillations during the first task (FA in FA-FP pair). Thus, the color reflects the power of alpha in reference to the given contrast; same color across brain areas means that the entire network is modulated by the task contrast in the same way and different color e.g in left vs right hemisphere means that in the two hemispheres or areas the power of alpha oscillation modulates to opposite direction in reference to the task contrast. The averages of FA-FP and FA-AT contrasts (in Figs 3A and 3B) appeared quite similar, both showing pattern of increased activation in the left and decreased activation in the right. Average of FP-AT contrast (in Fig 3C) shows a bilaterally symmetric pattern, and is somewhat weaker. Spatial contrast maps of FA-FP contrast for each individual participant are included in supplementary material (Figs A.2-A.5).

**Fig 3.**
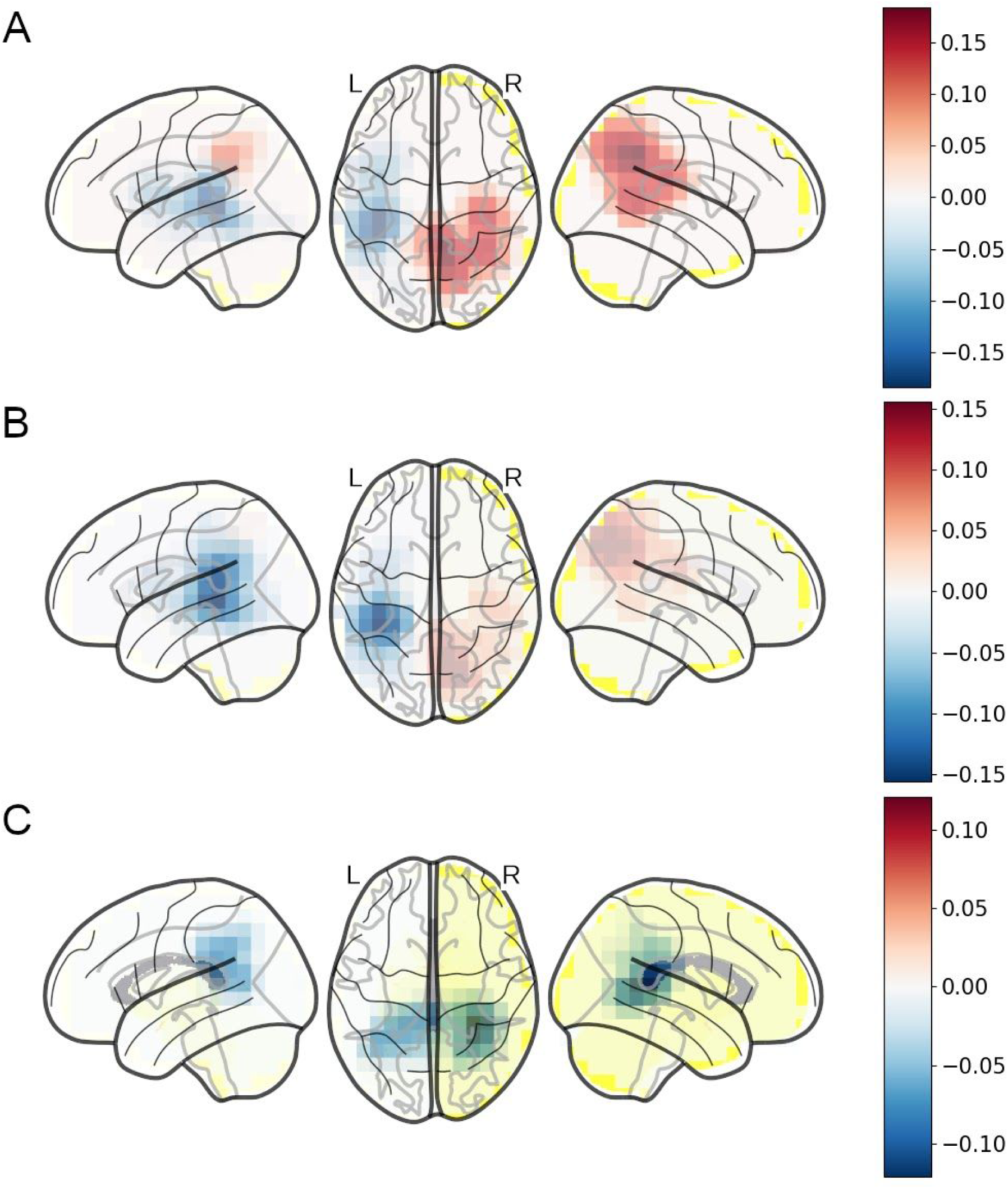
Averages computed from spatial contrast maps over 28 participants. The FA-FP contrast (A) shows a pattern of laterally opposite activations, which is quite similar to FA-AT (B). The contrast FP-AT in (C) shows a bilateral pattern.

### 3.3. PCA on spatial contrast maps

We ran PCA on the contrast maps to find most relevant subspaces in all three task pairs. The mixing weights for the first principal component in all task pairs are shown in Fig 4. Figs 4A and 4B illustrate the weights for FA-FP and FA-AT contrasts, respectively. They both show a component that captures a bilaterally symmetric brain network in parietal areas which increases/decreases in amplitude with respect to the contrast. Thus, in addition to the similarity of means shown in Fig 3, FA-FP and FA-AT contrasts seem to be similar in their correlation structure. Fig 4C shows a slightly more posterior pattern for the FP-AT contrast.

**Fig 4.**
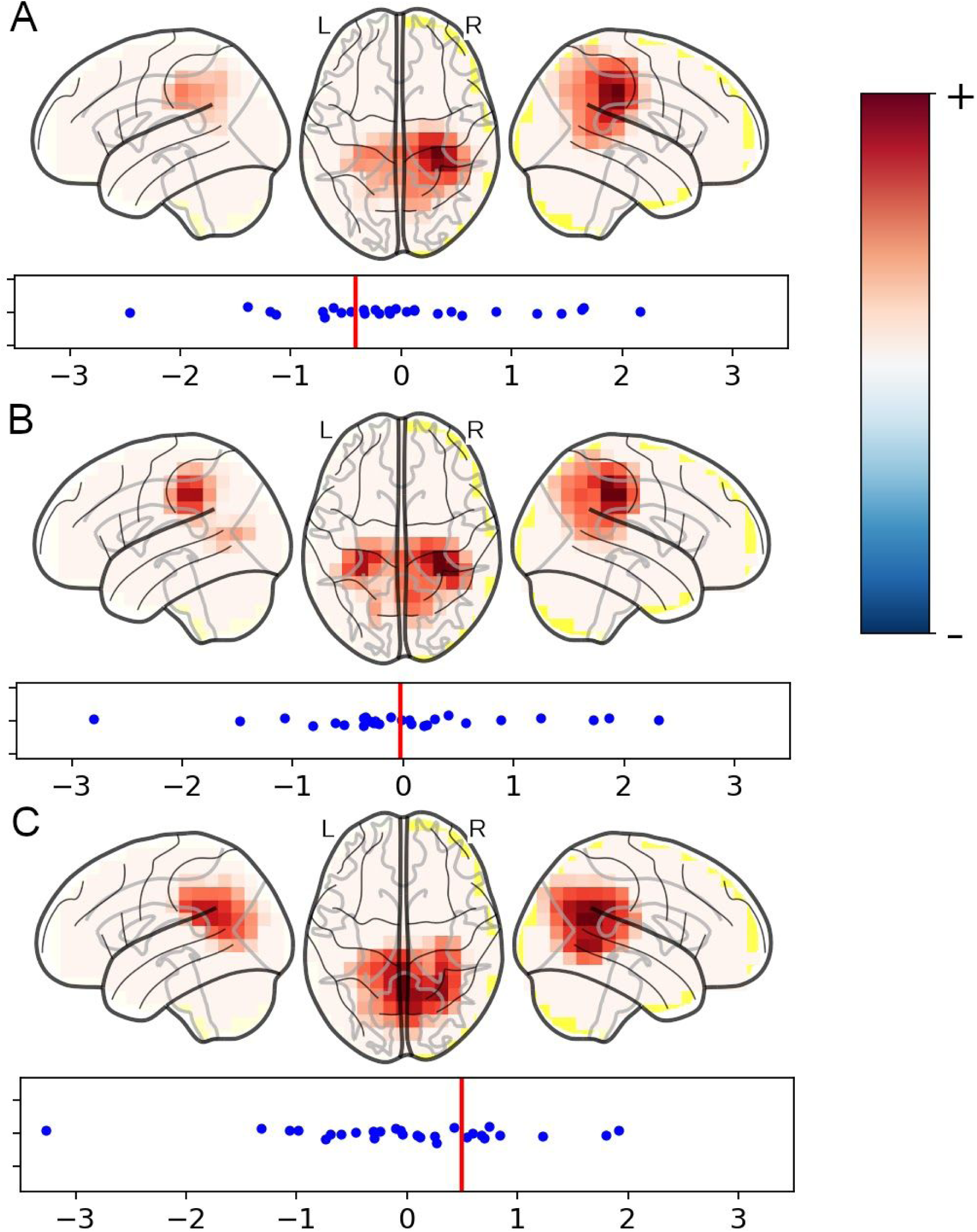
Mixing weights of the first PCA component in FA-FP (A), FA-AT (B) and FP-AT (C) contrasts, along with the component scores.

Fig 4 also includes the distributions of the component scores, that is, values for each individual contrast map after projecting them to principal axis using PCA. The vertical line shows the value of hypothetical contrast map containing only zeros, and basically reflects the effect of removing the mean before analysing covariances. It can be interpreted as a score for a participant who shows no difference between the tasks, and thus as a cutoff value where the difference between tasks changes direction. For example, because the average in Fig 3A shows positive value in the right parietal area, and because the mixing weights in Fig 4A are slightly stronger on the right, most of the participants are to the right of the vertical line in the component score plot. For the second component, the vertical line moves closer to the mean as the mixing weights are slightly more symmetrical, and the right parietal area of the mean in Fig 3B is somewhat weaker. Overall, even after taking the effect of mean into consideration, participants seem to exhibit quite variable patterns of activation, where even the signs of the effects can be opposite to each other.

The principal components, even with relatively low number of samples, were robust to individual participant omission, that is, leaving any of the participants out resulted always in a very similar set of components, with cluster quality index of around 0.8 for the first component of each pair. A dendrogram showing the results of clustering for FA-FP pair is included in supplementary material (Fig A.7). All three components for FA-FP pair are shown in Fig A.6. Components in other two task pairs were similarly robust. The first principal component in all task pairs explained substantially more variance than the other components (65% for FA-AT contrast, 65% for FA-FP contrast and 80% for FP-AT contrast), and was thus the only one kept for further analysis.

### 3.4. ICA on spatial contrast maps

As the first principal component explained most of the variance, the analysis above focused on using only it. Next, a more fine-grained analysis was done by taking more principal components, and in particular, doing ICA in the subspace spanned by the first principal components. For the FA-FP and FA-AT pairs, one independent component (see Fig A.8C) was similar to the first principal component, and behaved similarly in relation to subsequent regression. Another component (see Fig A.8A) had no connection to the BIA-component, but seemed to capture the amount of laterality that was seen in Figs 3A and 3B. The laterality seemed to be statistically significant (*p* = 0.001), that is, on average, participants exhibit increase of alpha oscillations in the right hemisphere and a decrease of alpha oscillations in the left hemisphere. These results suggest that the ICA decomposition (after averaging and contrasting the data) can be a viable complement to the PCA decomposition, and bring out relevant and meaningful aspects of the data. As it uses more information, however, it might be more subject to fitting to noise, and the robustness of components should be carefully analysed. Using the first principal component may be recommendable in the case of a small sample size, as was our case here.

### 3.5. Association of behavioral component to spatial activation maps

As a preliminary step, we investigated whether the tasks themselves were associated with the BIA-component computed from the questionnaire data. The component scores from the first principal component are often close to the means computed from each of the samples (participants in our case) separately. Thus for this initial investigation we took the mean of the spatial activation map for each participant and each task, and regressed the means with the BIA-component scores. This resulted in no correlation (*p* > 0.3) in all cases.

### 3.6. Association of behavioral component to spatial contrast maps

We next regressed the first principal component of contrast maps with the BIA-component for all three task contrasts (see Fig 5 for scatter plots.) We found that the FA-FP and FA-AT pairs are similarly correlated (*r* = 0.45, *p* = 0.015 for FA-FP, and *r* = 0.42, *p* = 0.024 for FA-AT pair) with the behavioral characteristics, and found no connection for the FP-AT pair (*r* = 0.06, *p* = 0.74).

**Fig 5.**
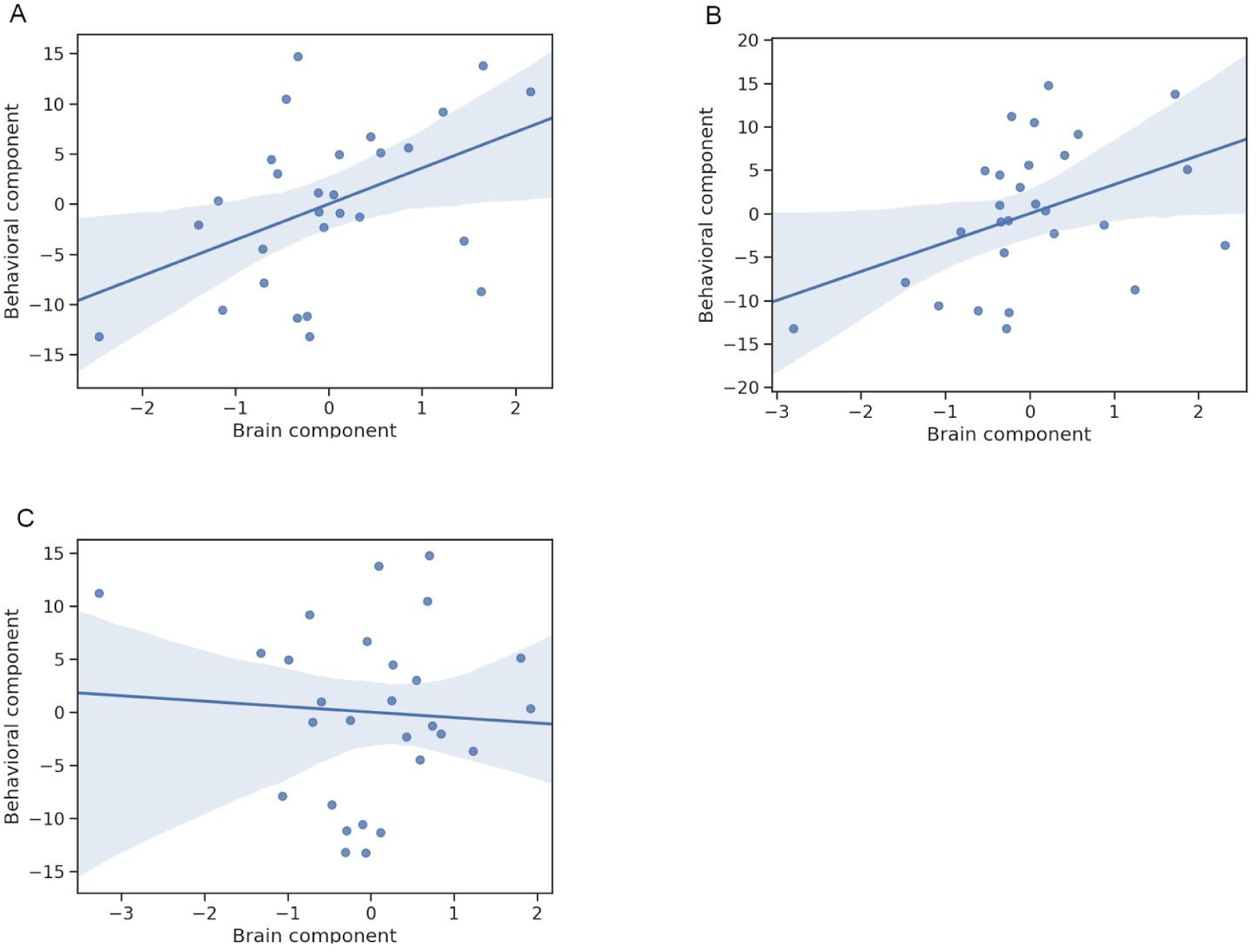
Scatter plots with regression lines for FA-FP (A), FA-AT (B) and FP-AT (C) contrasts.

In short, bilateral parietal network (see Figs 4A and 4B) is modulated by behavioral inhibition in such a way, that the more inhibited or anxious the participant is, the stronger the oscillations in mind wandering states are compared to focused attention. As the distributions of individual participants relative to principal components in Figs 4A and 4B show, the difference between states can be both positive or negative. Thus for the less inhibited participants, oscillations of mind wandering can be weaker than oscillations of focused attention, and for the more inhibited participants, oscillations of mind wandering can be stronger than oscillations of focused attention.

## 4. Discussion

Using data collected from continuous tasks of focused attention (FA), future planning (FP) and anxious thoughts (AT), we computed spatial activation maps for each task and participant, by averaging over respective time points. For each participant, we then subtracted condition-specific spatial activation maps from each other to get spatial contrast maps. Using PCA, we found distinctive subspaces along which the contrast maps of individual participants distributed, with notable variation across participants in the relative power across conditions in parietal regions. In particular, regressing the first principal component of spatial contrast maps in FA-FP and FA-AT contrasts with the trait of behavioral inhibition, anxiousness and depression, as measured by BIS, BAI and BDI questionnaires, revealed association in the parietal areas.

The usual approach in the analysis of trigger-free conditions with continuous electrophysiological data is to decompose the signal first by unsupervised methods such as ICA or PCA. In multi-participant context, concatenation methods are often used due to their ease of use. For example, in temporal concatenation, one first concatenates data from all participants in temporal dimension, then decomposes the concatenated signal, and then splits the decomposed concatenated signal to participant-specific decomposed signals. This way, it is straightforward to compare the values each component takes for each participant (Calhoun et al., 2009). As detailed in the introduction, this approach has drawbacks for investigating individual differences for at least two reasons. First, concatenation methods assume consistent mixing, i.e same spatial filter in temporal concatenation, across participants, which can be violated because of measurement errors or differences between brains. In addition, also individual-specific variation is present in the concatenated data, and thus some components may be based on variation present only in a small subset of participants, making it difficult to interpret values for participants that had negligible contribution. Some of the matter has been investigated in a simulation-based study (Allen et al., 2012), which highlights that concatenation methods are quite robust when differences between sources in different brains are small, but the quality deteriorates quickly when differences become larger. Specifically, it is noted that amplitude of component for single participant is correlated with how well the component is estimated for that participant. Second, due to many different sources of variance, such as noise, variance within- and between-tasks, and variance within- and between-participants, the decomposition usually results in a large number of components. Having a small number of components is good both for practical reasons, as it can be time-consuming to go through all components to find the useful ones (Hyvärinen et al., 2010), and for statistical reasons: there has been a lot of discussion recently on the low repeatability of psychological research due to questionable analysis practices (Simmons et al., 2011).

In the approach we took, the first problem is largely bypassed by averaging out the within-participant variance. This removes information that can bias the decomposition. There is, however, an inherent tradeoff in the process: we lose some intricate details of the brain activation to win in interpretability of the data in the special case of investigating individual differences. The second key feature of our approach is the contrasting of task averages within participants. Use of task contrast maps in the decomposition removes the variation within and between participants that is not related to the matter of interest. These two analysis steps combined thus leave only few but meaningful components for further analysis, and thus mitigate the second problem of having too many components.

The stability of principal components was evaluated using leave-one-participant-out procedure, similar to icasso (Himberg et al., 2004). The presence of individual participants had little effect on the estimated components. Especially the first component, which was the one used in regression analysis, had a very isolated cluster, indicating that it is robustly estimated. In all task contrasts, the first component explained substantially more variance than other components, and was for this reason the only component used in the later analysis. This resulted in a particularly parsimonious presentation and simple interpretation, where a single spatial pattern is used to explain the variability between participants.

We additionally decomposed the data with ICA to three components, for the FA-FP contrast. In terms of explained variance, this resulted in a more balanced set of two components, and one one component with negligible contribution. When comparing the ICA- and PCA -based component characteristics, one of the independent components had very similar spatial weights as the first principal component. Thus, to find the association between behavioral characteristics and the contrast maps, ICA could have also been used. The other component had a pattern where opposite hemispheres increased and decreased, respectively, in activation. It was not correlated with the BIA-component, but might still be meaningful, capturing the interhemispheric differences related to the task contrast. However, using ICA with such small sample sizes (less than 100) is quite risky since ICA usually needs much more data to give robust results.

Interestingly, the component scores and even the individual contrast maps (see Figs A.2-A.5) reveal a lot of variability across individuals. Some participants showed even the exact opposite patterns of activation to each other. This highlights the remarkable individual variability in the continuous MEG activation underlying these task conditions. It also explains why building group-level classifiers for the purpose of brain-computer interfaces is difficult with this type of data, as we attempted using these same recordings in a previous study (Zhigalov et al., 2019). More specifically, the cortical networks engaged during focused attention vs. mind wandering seem to show distinctive characteristics at individual level especially in bilateral parietal cortex. Importantly, by using linear regression to associate the first principal component of spatial contrast maps of FA-AT and FA-FP to behavioral inhibition – approach -component, we found them positively associated. Simply put, the higher the difference in alpha amplitude between mind wandering vs. focused attention, the higher the tendency for behavioral inhibition, evaluated using standardized questionnaires.

Behavioral inhibition was not correlated with the original spatial activation maps. This may stem from task-unrelated inter-individual variability in the MEG data. Therefore, we do not know if only one of the contrasted tasks was modulated by the behavioral characteristics, or if they both contributed to the observed association. If we assume that the participants have their brain activated in focused attention task in a similar fashion, and that increase in alpha oscillations signifies inhibition of brain networks, the result can be interpreted as more anxious or behaviorally inhibited participants showing less activation (or more inhibition) in parietal areas during mind wandering.

It is not surprising that the individual variability in behavioral tendency for inhibition and anxiety is associated particularly with activation in parietal areas. Bilateral parietal areas have originally been thought of as ‘association areas’ linking modality specific information e.g. from auditory and visual areas. However, beyond pure associative function, parietal regions have been recently linked especially with prioritizing the focus of attention and cognitive control (Bisley and Goldberg, 2010; Sapountzis et al., 2018). As our task conditions required controlling of attention, it is tempting to think that our results reflect activation of the frontoparietal control network, that has nodes located in the bilateral parietal areas (Cole et al., 2014). Frontoparietal control network has been suggested to support flexible switching between default and dorsal attentional network, and consequently the focusing of attention to internal, autobiographical information vs. external cognition (Smallwood et al., 2012; Spreng et al., 2010). Our tasks required focusing of attention either on ongoing sensory experience of one’s own body (breathing), or on visually triggered autobiographical thoughts (mind wandering). The results could indicate that inhibition (higher alpha) of the cognitive control/attentional resources during mind wandering are linked with tendency for anxiety and behavioral inhibition. Interestingly, in line with this interpretation, increased activity in the control network has been observed after successful psychotherapeutic treatments of depression and anxiety disorders. (Clark & Beck, 2010). However, the linkage to frontoparietal control network is still very speculative, as we did not see frontal areas sensitive to the studied contrast.

There are some limitations in the conducted study. The accuracy of inverse transform depends on the model of the brain, which usually is based on individual MRI images. For this study, however, we did not have MRI images for all participants, and instead, used a default template head model from freesurfer-package, to which the digitized head was aligned. Because the real shape of individual heads can vary, the accuracy of the inverse transform is slightly reduced, which must be kept in mind when interpreting the results. There are also limitations to the wider use of the methodological approach. It can only be applied to datasets that contain multiple continuous tasks, and is not applicable for example if only resting state recordings are available. It is also reasonable only when the interest lies in the individual differences, and not in common intricate activations. It is also worth noting that activation patterns shared by both tasks, which are lost in the subtraction, may still be significant determinants of individual variation.

In conclusion, this study explored the use of PCA of spatial contrast maps as a way to investigate individual differences. The ability of the method to extract behaviorally relevant, participant-specific characteristics of brain activity with no need for post-hoc component selection was demonstrated using a dataset containing continuous focused attention and simulated mind wandering states. Furthermore, the association of behavioral inhibition to the activation levels of parietal areas makes sense in the light of the previous literature giving evidence for the validity of the method. As the exploration of individual differences through neuroimaging methods has been an increasing trend in neuroimaging studies, we see a demand for methodology that can specifically target them in the context of continuous and more naturalistic settings. The validity of the method in wider applications for neuroscience should be confirmed and replicated in future studies.

## Acknowledgements

This work was supported by the Academy of Finland [grant number 295075], by CIFAR (A.H.) and by the Doctoral School of the Faculty of Information Technology of the University of Jyväskylä (E.H.). We are grateful for Iikka Pietilä for discussions, and help in the data recording phase.

## Appendix A: Supplementary figures

**Fig A.1.**
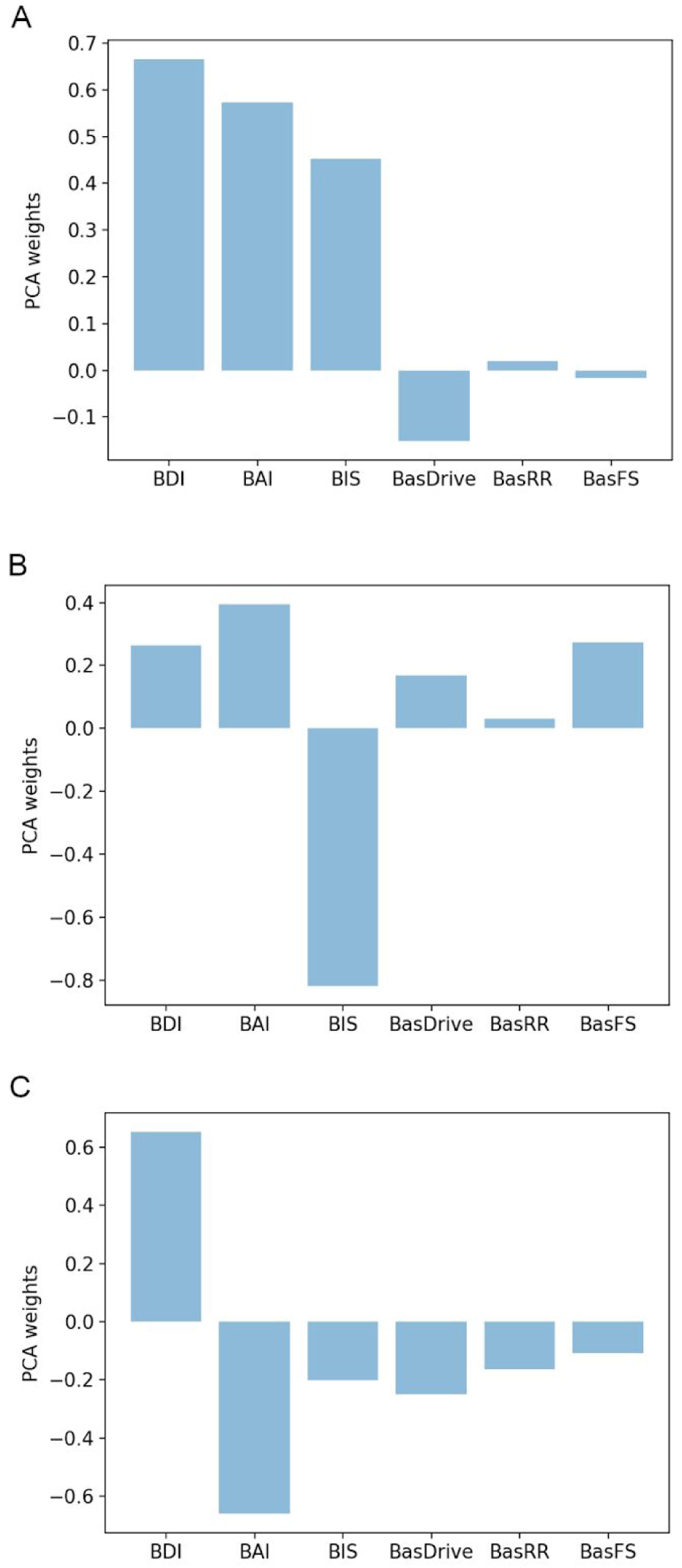
Mixing weights for first three PCA components from questionnaire data.

**Fig A.2.**
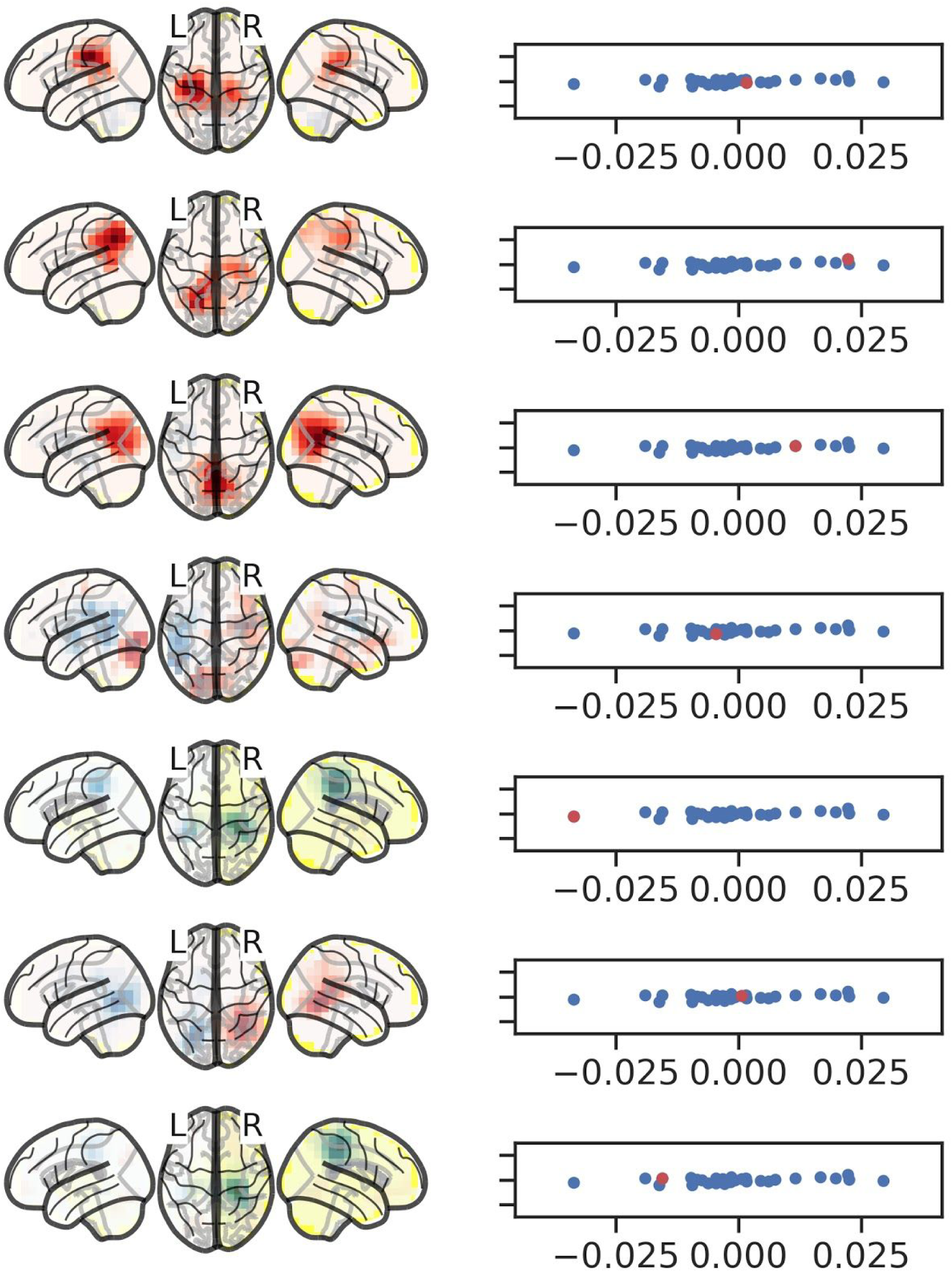
Spatial contrast maps (FA-FP) for participants 01-07. The first column contains spatial contrast maps for each participant, and the second column shows their projection to first principal axis (red) compared to other participants (blue).

**Fig A.3.**
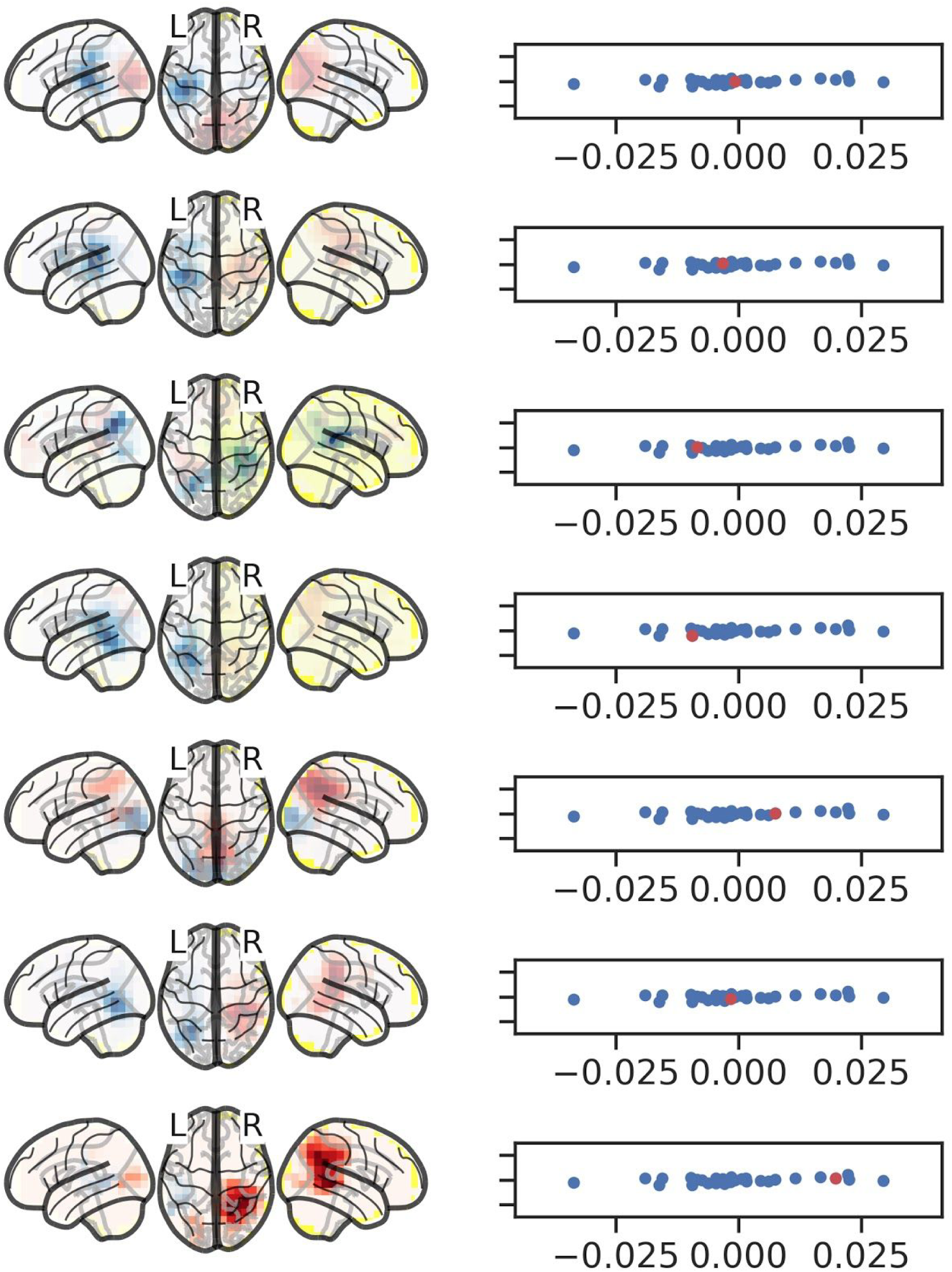
Spatial contrast maps (FA-FP) for participants 08-14.

**Fig A.4.**
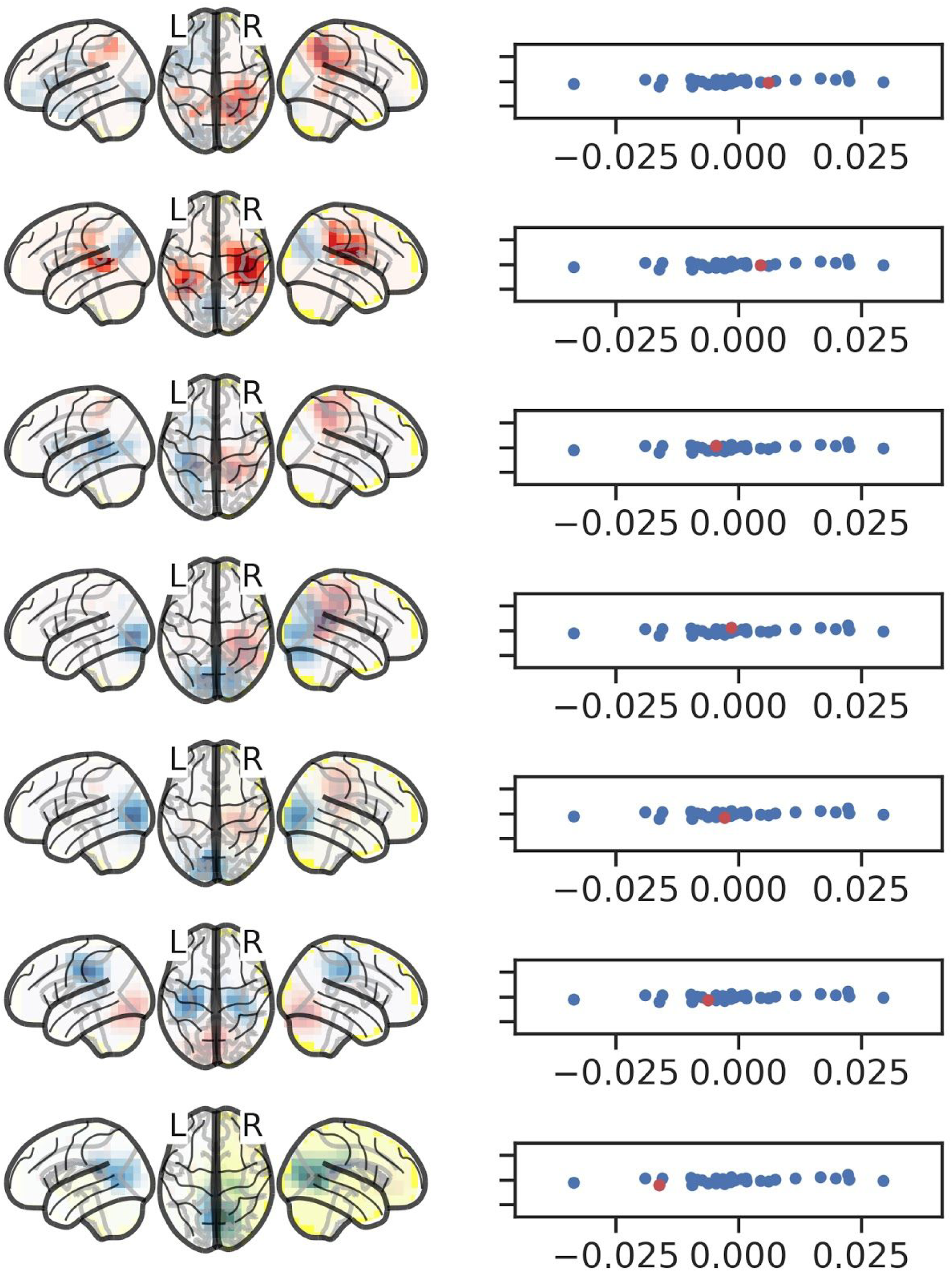
Spatial contrast maps (FA-FP) for participants 15-21.

**Fig A.5.**
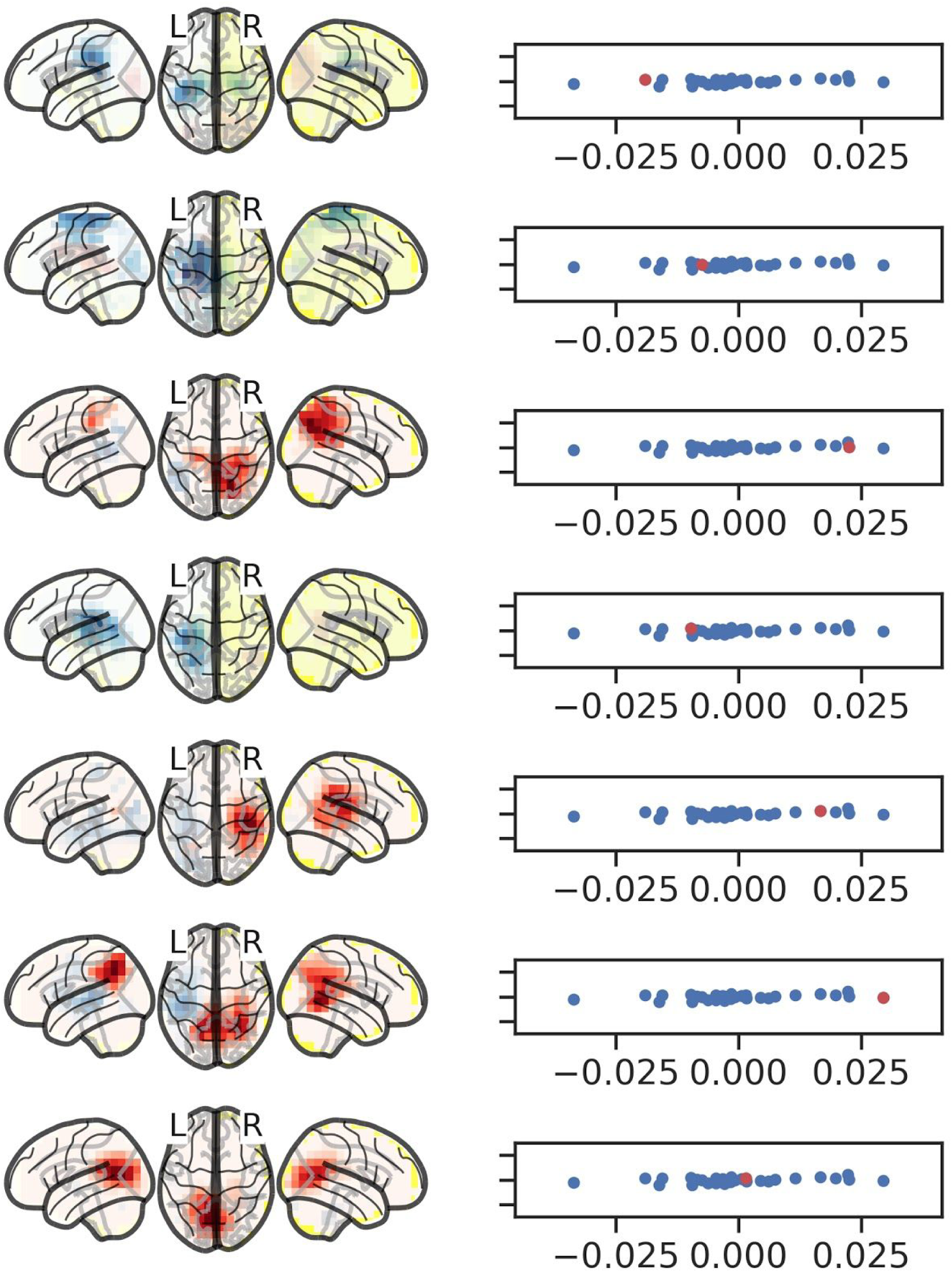
Spatial contrast maps (FA-FP) for participants 22-28.

**Fig A.6.**
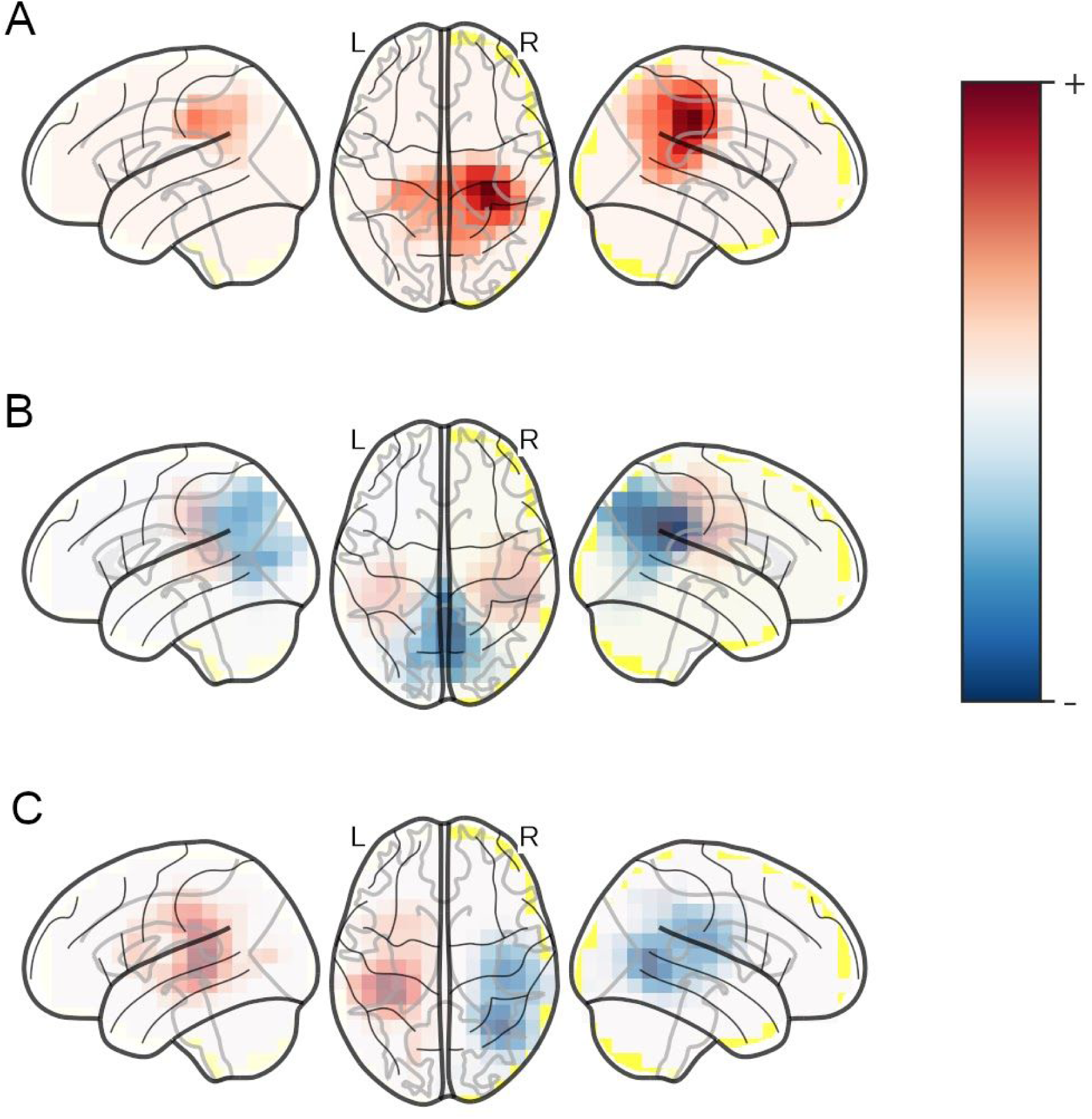
Mixing weights for first three PCA components computed from spatial contrast maps (FA-FP) from 28 participants.

**Fig A.7.**
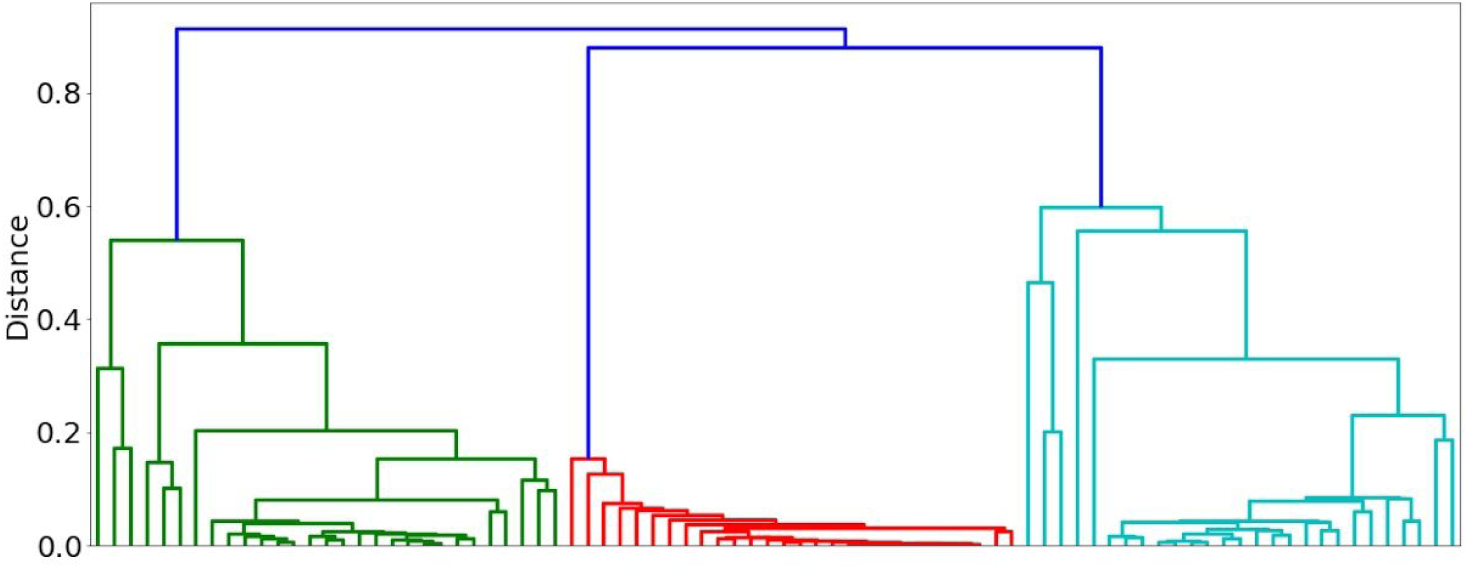
Dendrogram of the first three PCA components clustered using hierarchical clustering. Every intersection of vertical lines and the bottom of the figure represents a single component in a single run. The longer the vertical blue line is at the top, the more isolated the cluster is.

**Fig A.8.**
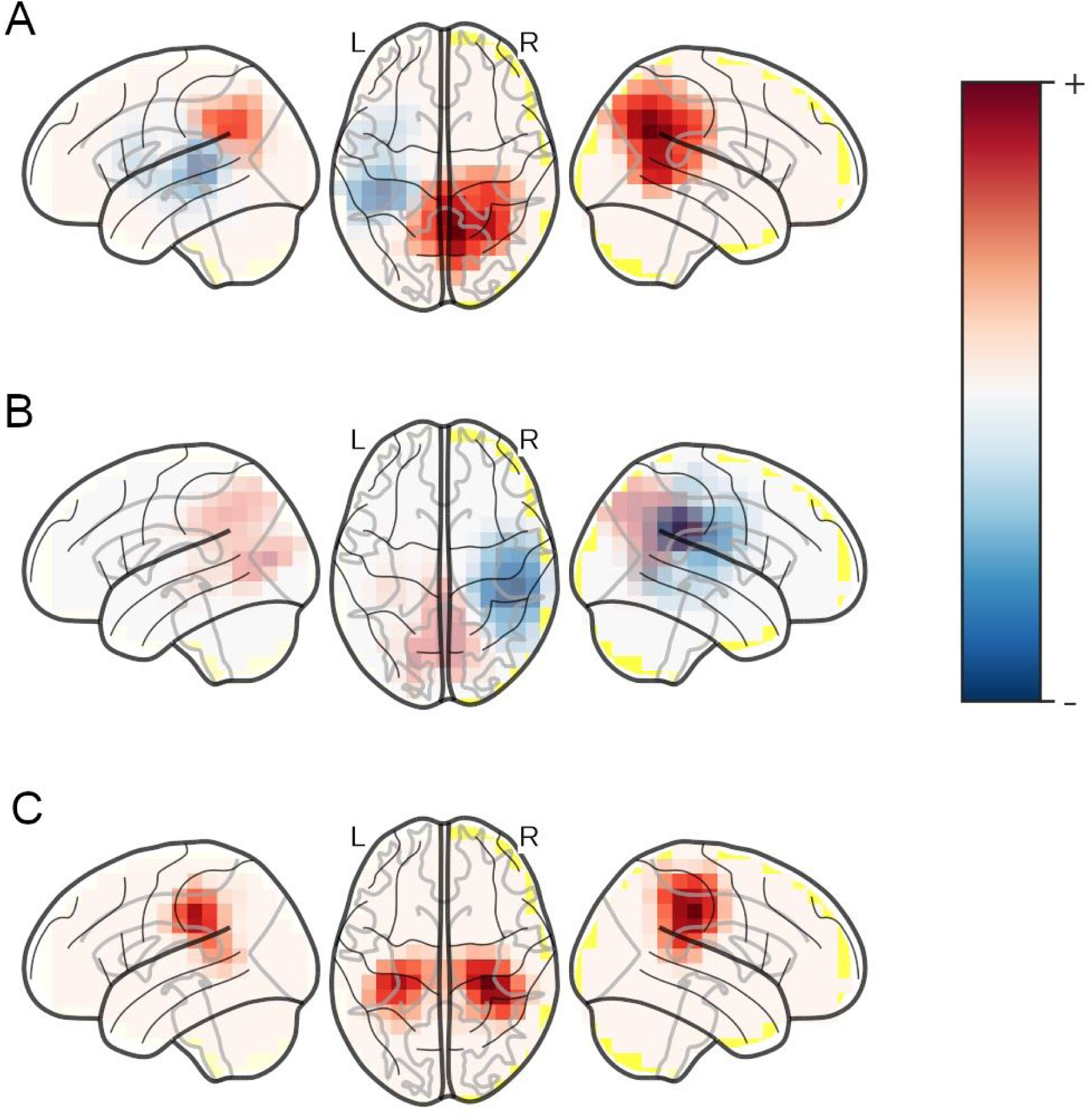
Mixing weights for the three ICA components computed after projecting the spatial contrast maps (FA-FP) to first three principal axes.

